# Chikungunya virus susceptibility increases in the absence of *mk2b* and *mk3* genes in zebrafish model

**DOI:** 10.1101/2025.03.24.644917

**Authors:** Supriya Suman Keshry, Usharani Nayak, Prabhudutta Mamidi, Suryasikha Mohanty, Udvas Ghorai, Rajeeb. K. Swain, Soma Chattopadhyay

## Abstract

Chikungunya virus (CHIKV) infection imposes a significant socio-economic burden due to the absence of effective antiviral treatments and vaccines. Host factors are essential for the CHIKV life cycle, making them promising targets for antiviral therapy. Previous studies have identified Mitogen-activated protein kinase activated protein kinase 2 (MK2) and Mitogen-activated protein kinase activated protein kinase 3 (MK3) as key host factors in CHIKV infection; however, their role in an animal model remain unclear. This study highlights the critical roles of MK2 and MK3 host factors in an animal model following CHIKV infection. To investigate their functions, *mk2b (mk2b-/-)*, *mk3 (mk3-/-)*, and *mk2b-mk3 (mk2b-/-mk3-/-)* double knockout zebrafish were generated using the CRISPR-Cas9 technique. A significant high viral titer of CHIKV was observed in case of all knockout groups compared to the wild-type (WT) control using plaque assay, RT-qPCR and immunofluorescence assay. Among the knockout groups, *mk3-/-* displayed the highest susceptibility to CHIKV, followed by *mk2b-/-.* In contrast, the *mk2b-/-mk3-/-* double knockout exhibited the lowest susceptibility to CHIKV infection. Additionally, severe symptoms such as bent body, impaired response to physical stimuli, and increased mortality were most pronounced in *mk3-/-* larvae compared to other knockouts and the WT. The expression levels of *infɸ1* and *rsad2* were also elevated in all knockout groups during the early days of infection indicating higher interferon response in the absence of *mk2b* and *mk3* during CHIKV infection. In conclusion, this study confirms that the *mk2b* and *mk3* host proteins are essential in controlling the CHIKV infection in organism level and subsequently may contribute in designing antiviral therapeutics in future. Furthermore, the knockout model of *mk2b* and *mk3* in zebrafish could serve as a valuable tool for studying their roles in other viral infections.

**Author Summary:** CHIKV, transmitted by *Aedes aegypti* and *Aedes albopictus* mosquitoes, causes febrile illness and has spread across Africa, Asia, Europe, and America. Despite extensive research, effective antiviral drugs and vaccines are yet to be commercially available. This study examines the role of *mk2b* and *mk3* host factors following CHIKV infection. For this purpose, *mk2b* and *mk3* single as well as double knockout have been generated in zebrafish using the CRISPR-Cas-9 technique. Findings suggest that zebrafish exhibit high CHIKV susceptibility in the absence of *mk2b* and *mk3*, confirming its importance during infection. Moreover, these three knockout models could serve as a valuable platform for examine the role of *mk2b* and *mk3* in the presence of other viral infection.

## Introduction

Chikungunya virus (CHIKV), is a mosquito-borne pathogen that is classified as a member of the *Alphavirus* genus within the *Togaviridae* family (1). The genome is characterized by a positive sense single-stranded RNA molecule that measures 11.8 kb in length. It is spherical enveloped virus, where nucleocapsid is surrounded by the glycoprotein E1-E2 on the plasma lipid membrane. The genome of CHIKV is divided into two open reading frames (ORFs) along with a 5’ cap and 3’ poly A tail. ORF1 encodes the non-structural proteins (nsP1, nsP2, nsP3, and nsP4) that are necessary for virus replication, while ORF2 translates the structural proteins (capsid protein C, E1, E2, and two small peptides E3 and 6K) (2,3). CHIKV was first identified as a human pathogen that caused debilitating arthritic disease in 1952 in Tanzania (3). The pathogenesis of chikungunya fever includes some non-specific symptoms like headache, nausea, rashes on the skin, vomiting and fever followed by specific symptoms like polyarthralgia, myalgia, and gastrointestinal complaints (3,4).

As obligatory parasites, viruses depend on both host and viral proteins for their replication and without host factors, virus progression is inexecutable. The viruses apply different strategies to hijack the host factors, which eventually leads to inhibiting or modulating the signaling pathways (5–8). One of the pathways that is activated after different types of stress conditions, including virus infection is the p38 mitogen activated protein kinase (p38-MAPK) pathway. Mitogen-activated protein kinase activated protein kinase 2 (MK2) and Mitogen-activated protein kinase activated protein kinase 3 (MK3) are the major downstream proteins of this pathway, which are activated by various stress conditions (9,10). These proteins are structurally and functionally similar, and are considered as isozyme partners of each other. They share a high degree of amino acid homology, with MK2 being 75% identical to MK3. MK2 and MK3 regulate a variety of substrates, ultimately leading to several key functions, such as actin remodeling, cytokine production, cell cycle control, and chromatin remodeling. The N-terminal region of MK2 and MK3 contains a proline-rich area (10-40 aa) followed by a catalytic domain (64-324 aa), while the C-terminal region contains a nuclear export signal (NES) (356-365 aa), nuclear localization signal (NLS) (371-374 and 385-389 aa), and a p38 binding domain. The primary phosphorylation sites for these proteins are located at T222 and T334 (11).

The MK2 and MK3 proteins become activated following stress conditions, like infection with viruses. For instance, studies have shown that MK2 and MK3 host proteins aid the chikungunya virus in egressing out from the cells through the actin remodeling pathway. Silencing of these kinases using small interfering RNA or inhibitors both *in vitro* and *in vivo* has been demonstrated to reduce the viral load in the extracellular environment (12). Similarly, in the case of influenza virus, a significant reduction in viral titer has been observed in the cells deficient in MK2 and MK3 expression. Additionally, the knockout of MK2 and MK3 was found to affect influenza viral protein synthesis but not viral mRNA levels (13), and similar findings have been reported for the Hepatitis-C virus (14). To explore the function of these two host proteins in greater details, the development of knockout animal models is a valuable approach.

Zebrafish (*Danio rerio*) is a useful animal model for studying the role of genes under physiological conditions (15). It serves as a valuable experimental system to investigate infections for a wide variety of pathogens due to their genetic and physiological similarities with higher vertebrates, including humans. Zebrafish larvae are well-known for studying host-pathogen interactions because of their optical transparency, genetic manipulability, and translational potential (15,16). The development of the zebrafish immune system is well understood, making the use of larvae ideal for investigating innate immunity (16,17). Due to the advantages of zebrafish in real-time observation, genetic editing, and high-throughput drug screening, it serves as an excellent technological platform for investigating host-virus interactions. Various viral infections have been standardized in zebrafish to study viral pathogenesis, virus-induced immune responses, involvement of host factors, and the evaluation of antiviral compounds (17–21). According to Palha et al. (20), CHIKV infects multiple organs of zebrafish, however ultimately persists in brain parenchyma and is cleared from the body through type-I interferon responses. Neutrophils are the primary population of leukocytes that produce interferon and play a key role in controlling chikungunya virus infection.

In this study, the roles of *mk2b* and *mk3* were explored following CHIKV infection in the zebrafish model. Three zebrafish knockout lines which include single knockout of *mk2b (mk2b-/-), mk3 (mk3-/-)*, and double knockout of *mk2b* and *mk3 (mk2b-/-mk3-/-)* have been developed to evaluate the importance of these host proteins during chikungunya virus infection.

## Results

### *mk2* and *mk3* of zebrafish share a high degree of gene and protein homology with that of human MK2 and MK3

The genes *mapkapk2a*, *mapkapk2b*, and *mapkapk3* of zebrafish were retrieved from the ensemble database (mapkapk2a: ENSDARG00000002552, mapkapk2b: ENSDARG00000018530, and mapkapk3: ENSDARG00000111612). According to the Zfin database, two *mapkapk2* paralogs (*mapkapk2a (mk2a)* and *mapkapk2b (mk2b)*) exist in zebrafish. *mk2a* is located in chromosome number 11 and its position is from 21,154,678 to 21,218,172 base pair. The chromosome number of *mk2b* gene is 8 and its position is 37,465,380-37,477,845 base pairs whereas *mk3* is encoded by chromosome number 11 in between the base pairs 34,3 22,176 to 34,367,076. The study by Holloway et al.(22) identified the importance of *mk2a* in epiboly. The *mk2a* mutants were not viable and hence, this gene was excluded from the current knockout study, despite its greater sequence similarity to human MK2 (Fig. 1A). The gene and protein similarity of MK2 between zebrafish and humans was found 70.87% and 74.78% respectively, whereas the similarity of gene and protein of MK3 between zebrafish and human was 75.75% and 74.4% respectively (Fig. 1A and 1B). The *mk2b* and *mk3* genes encode transcripts of 397 and 408 amino acids respectively. Exon 5 of these proteins was chosen as the target site for knockout generation due to its pivotal location within the catalytic domain. Furthermore, the essential phosphorylation site, along with the NES, NLS, and the p38 binding site, are all positioned beyond Exon 5, reinforcing its suitability as an optimal target site. (Fig. 1C). Moreover, the data of multiple sequence alignment also suggested that all the key phosphorylation sites are conserved between zebrafish and human MK2 and MK3 proteins (Fig. 1D and 1E). Therefore, the analysis indicates that *mk2b* and *mk3* proteins of zebrafish show a high degree of protein homology with that of human MK2 and MK3 and thus can be taken for further knockout study.

**Figure 1:**
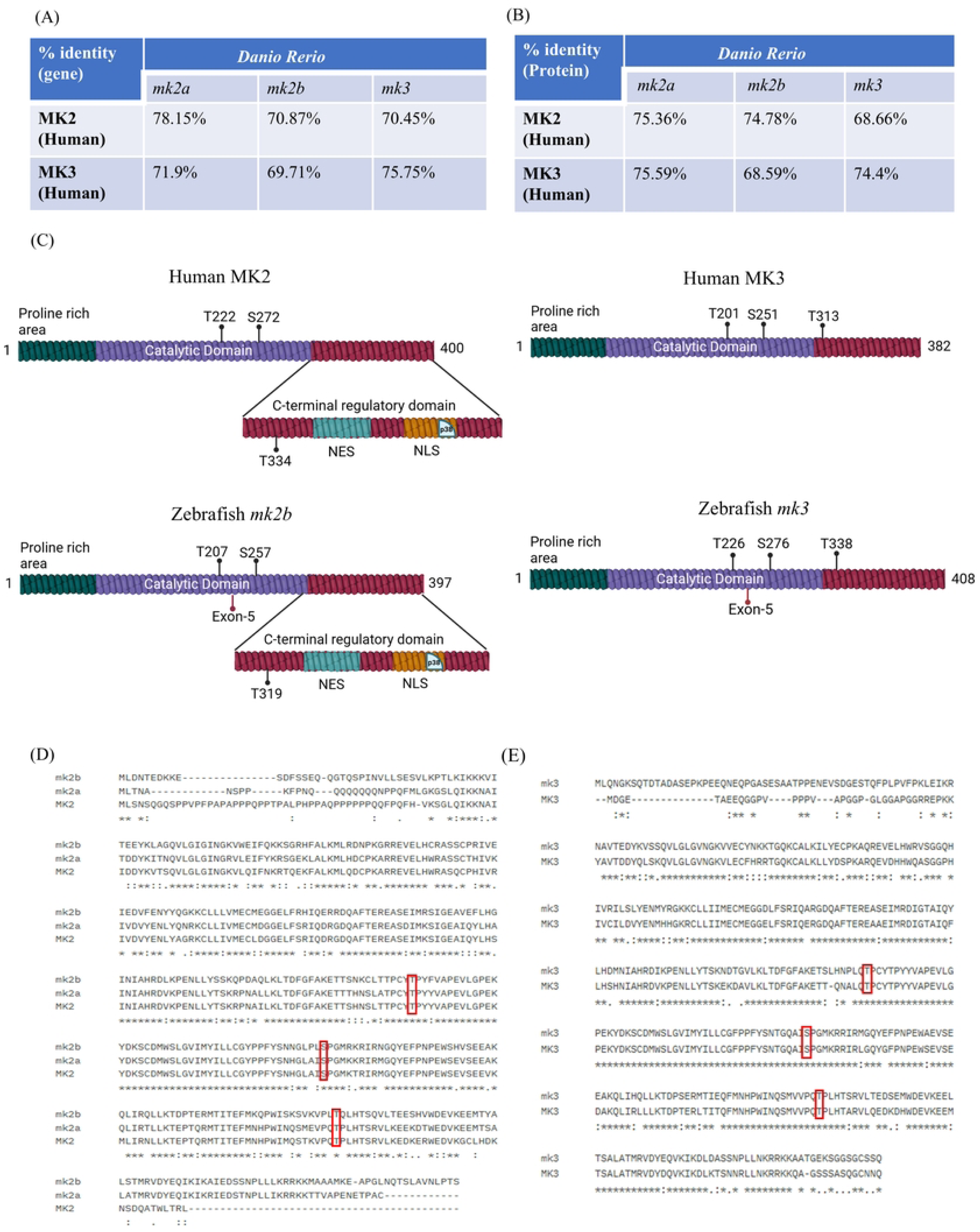
mk2 and mk3 of zebrafish share a high degree of gene and protein homology with that of human MK2 and MK3: In zebrafish, there are two isoforms of *mk2* - *mk2a* and *mk2b*. (A) Percent nucleotide identity of MK2 and MK3 genes between zebrafish and humans (B) Percent amino acid identity of MK2 and MK3 proteins between zebrafish and humans. (C) Different domains of MK2 and MK3 in humans and zebrafish, with the essential phosphorylation sites and position of exon-5 (target site for CRISPR knockout) in the domain. (D) Multiple sequence alignment of the zebrafish *mk2a*, *mk2b* and MK2 protein sequences of humans using the Clustal Omega tool. (E) Multiple sequence alignment of the zebrafish and human MK3 proteins using the Clustal Omega tool.

### Generation of *mk2b-/-, mk3-/-* and *mk2b-/-mk3-/-* zebrafish knockouts

To develop a single knockout of the *mk2b* and *mk3*, *mk2b* and *mk3* gRNAs along with the Cas9 mRNA were co-injected into the single-cell stage of zebrafish embryos separately and determined the mutation efficiency by genotyping. The process of genotyping involves genomic DNA isolation from the whole embryo or from the fin of adult fishes, high-resolution melt curve analysis (HRM) and heteroduplex (HD) analysis. The mutation efficiency of *mk2b* and *mk3* gRNAs were observed to be 80% and 90% respectively. For the generation of *mk2b* and *mk3* double knockout, both the gRNAs were microinjected in the same single cell stage embryo along with the Cas9 mRNA followed by genotyping. The microinjected larvae (F0) were grown for 3 months, and then checked for the positive mutants. The adult F0 mutants were crossed with wild types to get the F1 heterozygous. After genotyping, positive F1 fishes were sequenced to identify the type of mutation that was generated. Different types of mutations were identified at the target locus but only one mutation led to the frameshifting of the *mk2b* and *mk3* open reading frames, which created pre-mature termination codon (PTC) insertion. A comparison of chromatograms between wild-type and *mk2b* heterozygous mutant showed 26-nucleotides deletion at the targeted site of the *mk2b* gene (Fig. 2A). In case of the *mk3* heterozygous mutant, insertion of 26 nucleotides has been observed (Fig. 2B). Apart from these single knockouts, double knockout showed 26 nucleotides deletion and 20 nucleotides deletion in the target site of *mk2b* and *mk3* genes (Fig.2G) respectively, which led to shift in open reading frame and generation of PTC. These positive F1 heterozygous mutants of all three knockout lines (*mk2+/-, mk3+/- and mk2b+/-mk3+/-*) were selected to generate F2 homozygous fishes (*mk2-/-, mk3-/- and mk2b-/-mk3-/-*). Homozygous mutant fishes from all knockout groups were identified by the presence of band appearing either above or below the wild-type (WT) *mk2b* and *mk3* bands (Sup fig-1). To detect the decay of *mk2b* and *mk3* mRNA due to PTC insertion, WISH was performed using a DIG labelled antisense RNA probe (digoxigenin), and it was observed that there was almost no expression of *mk2b* and *mk3* mRNA in respective mutants, whereas WT embryos showed expected ubiquitous expression (Fig. 2C, 2E and 2H). The expressions of *mk2b* and *mk3* in all the knockouts were further quantified through RT-qPCR which showed reduced levels of *mk2b* and *mk3* in *mk2b-/-,* and in *mk3-/-* respectively and both the mRNAs were reduced in *mk2b-/-mk3-/-* double knockout (Fig. 2D, 2F and 2I). Together, the data confirmed that functional mutation of *mk2b* and *mk3* genes have been developed successfully in both single as well as double knockout sets.

**Figure 2:**
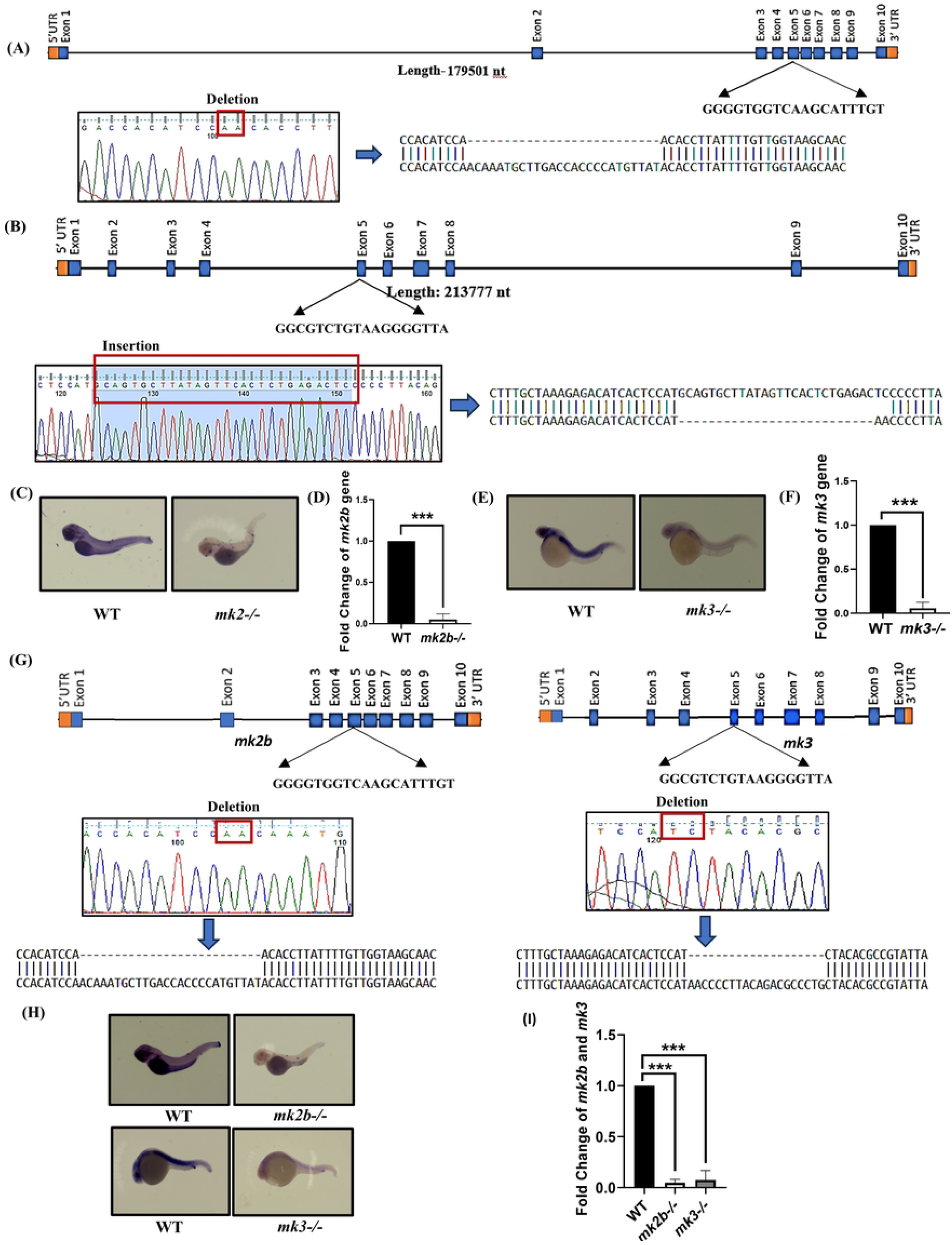
Generation of *mk2b-/-, mk3-/- and mk2b-/-mk3-/-* zebrafish knockouts: (A) A gRNA targeting exon-5 of mk2b was designed, synthesized and microinjected into the single cell stage zebrafish embryos, along with Cas9 mRNA. Chromatogram image indicating the type of mutation developed after introducing *mk2b* gRNA. (B) The gRNA of *mk3* was microinjected along with the Cas9 mRNA in the single cell stage of embryo. A chromatogram showing the mutation following the introduction of the *mk3* gRNA. (C) Microscopic images of zebrafish showing the expression of *mk2b* mRNA between WT and *mk2b*-/-. (D) Bar diagram showing the fold change of *mk2b* gene expression level. (D) Microscopic images of zebrafish indicating the expression of *mk3* mRNA between the WT and *mk3-/-.* (F) Bar diagram showing the fold change of *mk3* gene expression level. (G) gRNAs targeting the exon 5 regions of both the *mk2b* and *mk3* genes, were co-injected with the Cas9 mRNA into single-cell stage of zebrafish embryos. Chromatogram image representing the mutation of both the genes (*mk2b* and *mk3*) in the same mutated larvae. (H) Images of zebrafish larvae depicting the mRNA expression levels of *mk2b* and *mk3* between WT and *mk2b-/-mk3-/-* double knockout using ISH. (I) Bar diagram showing the fold change of *mk2b* and *mk3* gene expression in *mk2-/-mk3-/-* double knockout zebrafish larvae compared to WT. Data presented as mean ± SD. n=3; ***p=0.0003 and p=0.0008 were considered significant.

### Chikungunya viral titre increases significantly in the absence of *mk2b* gene in zebrafish

To assess the CHIKV infectivity in the absence of *mk2b* gene, approximately 250 CHIKV viral particles were microinjected to the common cardinal vein (CCV) in 3 days old larvae. A total of 15 larvae were collected daily for 5 days post infection (dpi), crushed and plaque assay was performed to assess the viral titer. A higher number of plaques were observed in *mk2b-/-* larvae on each day compared to the WT (Fig. 3B and C). The infectious viral particle load peaked on 2 dpi, exhibiting a 75.9% increase in viral titer compared to WT. This was followed by a decline in viral titer from 3 to 5 dpi, suggesting effective viral clearance in both WT and mutant larvae. The viral gene expression was also monitored using RT-qPCR, which indicated higher CHIKV-E1 gene expression in case of *mk2b-/-* larvae (Fig 3D). Infection of CHIKV was further confirmed by the presence of E2 protein in immunofluorescence assay in 2 dpi (Fig 3E). Altogether, the results suggest that *mk2b* is an important host protein that is associated with anti-CHIKV activity in zebrafish.

**Figure 3:**
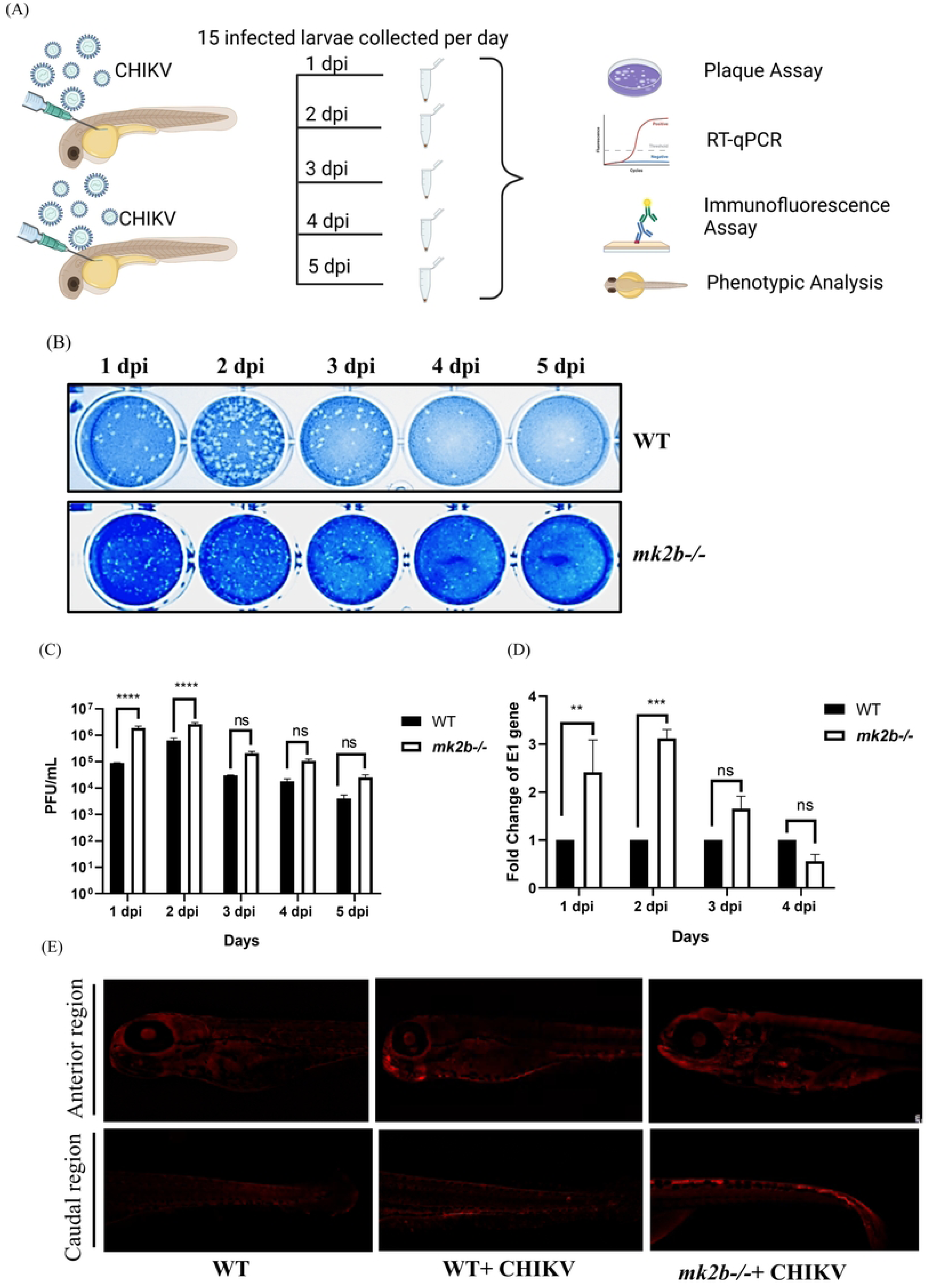
Chikungunya viral titre increases significantly in the absence of *mk2b* gene in zebrafish: 3 days old *mk2b-/-* larvae were microinjected with approximately 250 CHIKV viral particles intravenously (CCV). A total of 15 infected larvae were collected from day 1 to day 5 post-infection to assess the viral load in WT and *mk2b-/-* larvae. (A) Schematic diagram of virus injection and viral load detection by different methods. Created with BioRender.com. (B) Image of plaque assay plates showing viral load. (C) Bar graph representing the viral titer/mL in the *mk2b-/-* and WT zebrafish larvae. (D) Total RNA was isolated from WT and *mk2b-/-* zebrafish larvae after infection and the CHIKV-E1 gene was amplified by RT-qPCR. Bar diagram showing the fold changes of viral RNA. (E) Immunofluorescence assay depicting the CHIKV infection in the anterior and caudal region. The head is towards the left in all the pictures. Data of three independent experiments are shown as mean ± SD. ****p<0.0001 was considered significant.

### *mk3-/-* showed higher CHIKV titer compared to *mk2b-/-* and wild type zebrafish larvae

In order to examine the role of *mk3* during CHIKV infection, WT and *mk3-/-* zebrafish larvae were microinjected with approximately 250 CHIKV virus particles and were subjected to plaque assay, RT-qPCR and immunofluorescence assay as described above to detect the viral titer. A significant increase in viral load was observed in *mk3-/-* zebrafish larvae compared to the WT and *mk2b-/-* larvae (Fig. 4A and B), with a 90.6% increase in viral titer in 2 dpi. Similarly, viral gene expression was significantly higher in *mk3-/-* zebrafish (Fig. 4C). Further, CHIKV infection was confirmed by the presence of the E2 protein through immunofluorescence (Fig. 4D). Collectively, the data suggest that *mk3* is an essential host factor for controlling CHIKV infection in zebrafish.

**Figure 4:**
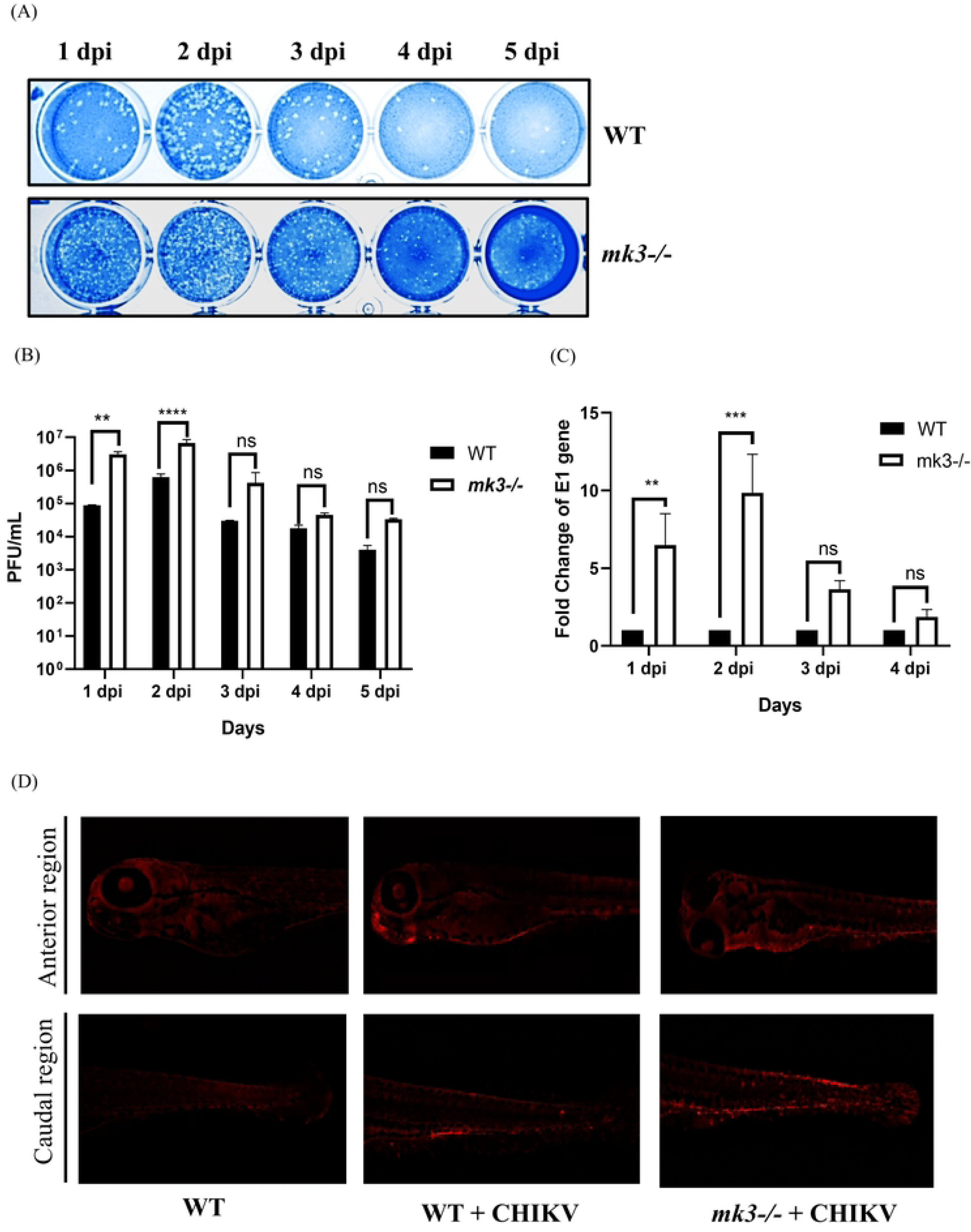
*mk3-/-* showed higher CHIKV titer compared to *mk2b-/-* and wild type zebrafish larvae: 3-day-old zebrafish larvae, produced from the *mk3-/-* parents, were intravenously microinjected with approximately 250 CHIKV viral particles. A total of 15 infected larvae were collected daily from day 1 to day 5 post infection to evaluate the viral load in both WT and *mk3-/-* larvae. (A) The image displays plaque assay plates indicating the viral load. (B) The bar graph illustrates the viral titer/mL in *mk3-/-* and WT zebrafish larvae. (C) Total RNA was isolated from WT and *mk3-/-* zebrafish larvae after infection and the CHIKV-E1 gene was amplified by RT-qPCR. Bar diagram showing the fold changes of viral RNA. (D) Immunofluorescence assay depicting the CHIKV infection in the anterior and caudal region. The head is towards the left in all the pictures. Data of three independent experiments are shown as mean ± SD. ****p<0.0001 and **p=0.0001 were considered significant.

### *mk2b-/-mk3-/-* double knockout zebrafish indicates less susceptibility to the CHIKV infection compared to the other knockouts

Since MK2 and MK3 are isozymes of each other, it is essential to understand their function during CHIKV infection when both the genes are impaired. Therefore, CHIKV infectivity was also assessed in the absence of both genes. To determine the CHIKV titer in *mk2b-/-mk3-/-* double knockout zebrafish, plaque assay was conducted as described above. The results revealed a significant increase (47.5%) in viral titer (Fig. 5A and B) in the *mk2b-/-mk3-/-* double knockout group compared to the WT control. However, the *mk2b-/-mk3-/-* double knockout exhibited lower susceptibility to CHIKV infection than other knockout groups. This elevated viral load was further validated by amplifying the CHIKV-E1 gene, which corroborated the plaque assay data by showing a marked increase in viral gene expression in the *mk2b-/-mk3-/-* double knockout group (Fig 5C). Additionally, immunofluorescence assays with CHIKV E2 protein confirmed successful CHIKV infection. (Fig 5D). Altogether, the data suggest that both *mk2b* and *mk3* genes are required to control the CHIKV infection.

**Figure 5:**
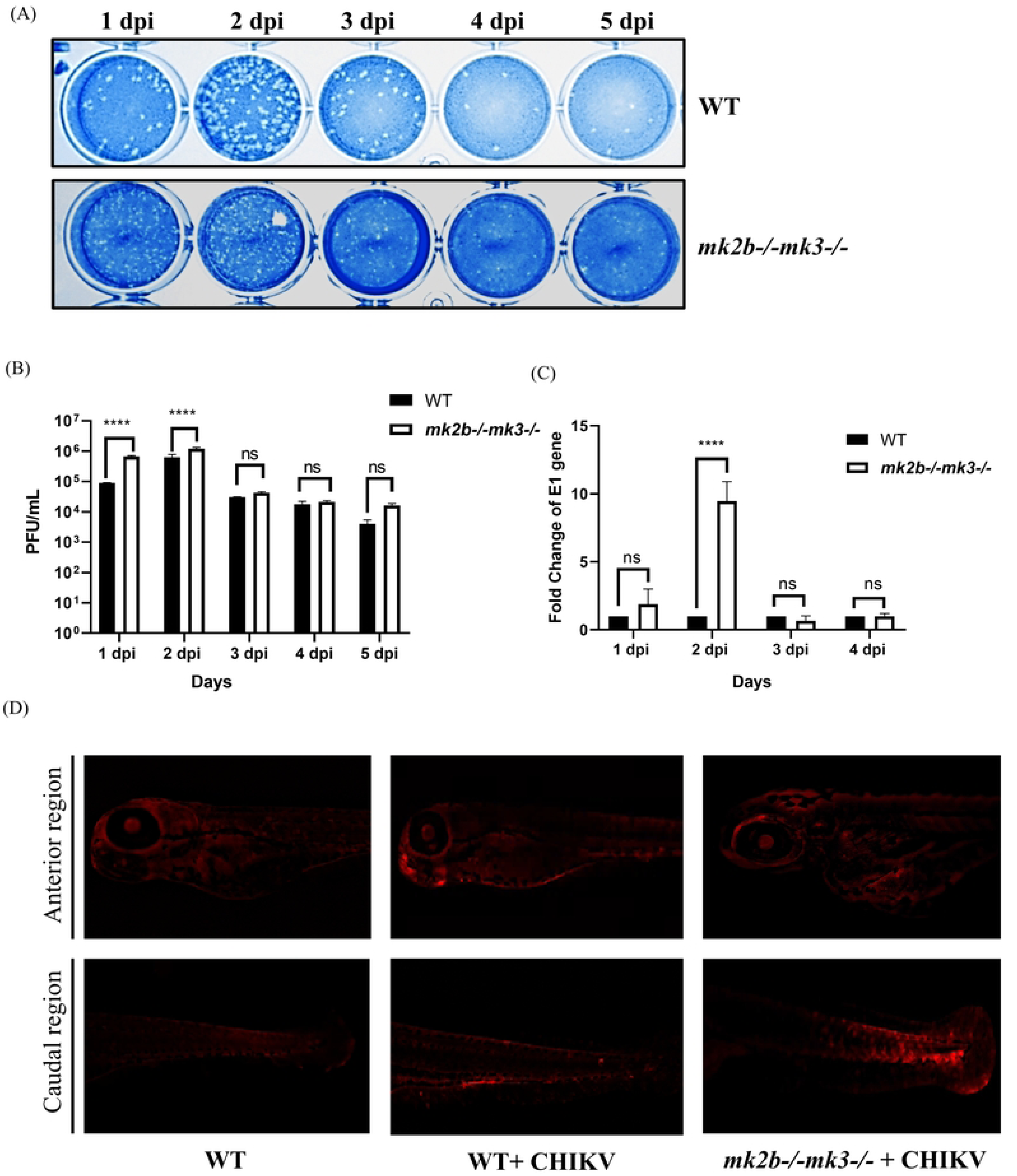
*mk2b-/-mk3-/-* double knockout zebrafish indicates less susceptibility to the CHIKV infection compared to the other knockouts: 3 days old *mk2b-/-mk3-/-* double knockout zebrafish larvae were microinjected with 250 CHIKV viral particles intravenously. Approximately15 infected larvae were collected from day 1 to day 5 to assess the viral load in WT and *mk2b-/-mk3-/-* double knockout larvae. (A) Image of plaque assay plates showing viral load. (B) Bar graph representing the viral titer/mL in the *mk2b-/-mk3-/-* double knockout and WT zebrafish larvae. (C) Bar diagram showing the fold change of CHIKV-E1 gene from infected sets of WT and *mk2b-/-mk3-/-* double knockout zebrafish larvae. (D) Immunofluorescence assay depicting the CHIKV infection in the anterior and caudal region. The head is towards the left in all pictures. The values of viral titer/mL and fold change of E1 gene were same for WT control in all the knockout condition (*mk2b-/-, mk3-/-* and *mk2b-/-mk3-/-*double knockout). Apart from this, Immunofluorescence imaging for WT and WT+CHIKV samples were kept common for all the knockout panels. Data of three independent experiments are shown as mean ± SD. ****p<0.0001 was considered significant.

### *mk3-/-* larvae show highest mortality and other symptoms after CHIKV infection

To compare disease pathogenesis of CHIKV in zebrafish, 250 CHIKV particles were microinjected intravenously (CCV) into WT and knockout larvae (3dpf). For clinical scoring, infected zebrafish were monitored daily from the 24 hours of infection (Day 1) to day 7. Disease scores were set based on symptom severity, which was recorded by increasing the viral titer through microinjection in WT zebrafish. Symptoms such as bent body, impaired response to physical stimuli, and mortality were observed in a higher number of larvae which correlated with high viral titers (Sup fig 2). Due to their severity, these symptoms were assigned higher clinical scores. These three symptoms were more prevalent in the case of *mk2b-/-* and *mk3-/-* larvae (Fig 6A, B and C). In *mk2b-/-* larvae, approx. 33% infected larvae died at the end of day 3, whereas *mk3-/-* infected larvae exhibited a mortality rate of 54%. In contrast, no mortality was observed in either the WT or *mk2b⁻/⁻ mk3⁻/⁻* double knockout group until Day 3 (Fig. 6A). A larger proportion of WT larvae displayed edema, abnormal heart rate, and irregular blood flow, whereas only a few showed bent body phenotypes (Fig. 6C, D and E). The symptoms of *mk2b⁻/⁻ mk3⁻/⁻* double knockout larvae appeared similar to those of WT. 0However, larvae displaying any symptom finally died between Day 4 to Day 7. Collectively, these findings suggested that virus infected *mk3-/-* larvae exhibited greater clinical severity compared to other knockout groups and WT zebrafish larvae.

**Figure 6:**
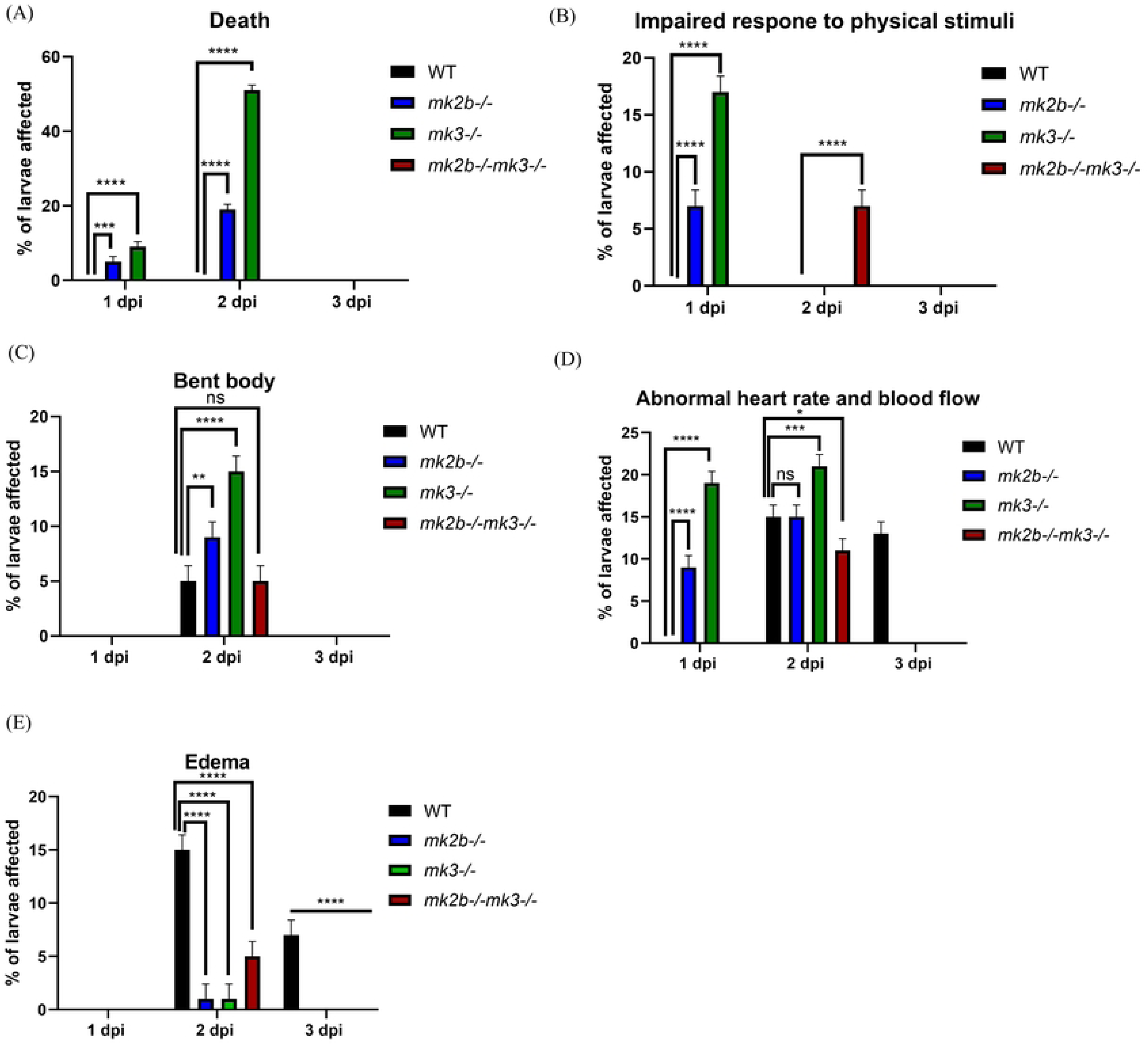
*mk3-/-* larvae show highest mortality and other symptoms after CHIKV infection: Both WT and knockout larvae (including *mk2b-/-*, *mk3-/-*, and *mk2b-/-mk3-/-*) were exposed to approximately 250 viral particles and monitored over three-days. Panels (A, B, C, and D) illustrated the percent of larvae showing the severity of symptoms in WT, *mk2b-/-, mk3-/-,* and *mk2b-/-mk3-/-* zebrafish larvae, respectively, with the following scale: 1 = Edema, 2 = Abnormal blood flow and heart rate, 3 = Bent body, 4 = Impaired response to physical stimuli, 5 = Death. Data presented as mean ±SD. n=3; **** indicates p<0.0001, *** p=0.0003, ** p=0.0018 and * p=0.0139 were considered significant.

### Expression levels of interferon and interferon induced gene were upregulated after CHIKV infection

Host triggers interferon production as the first stress response to viral infection which leads to activation of interferon induced genes and other proinflammatory cytokines production (23)). Mitogen-activated protein kinase (MAPK) is one of the pathways which regulates proinflammatory cytokines, with MK2 and MK3 serving as key downstream regulators. In order to understand the expression level of interferon (*ifnɸ1)* and interferon induced gene (rssad2) in the absence of MK2 and MK3, different knockouts (*mk2b-/-, mk3-/-* and *mk2b-/- mk3-/-)* and WT zebrafish larvae were microinjected with equal titer of CHIKV and its levels were quantified by RT-qPCR. Knockout larvae exhibited higher expression level of *ifnɸ1* on day 1 which gradually decreased by day3 (Fig 7A). On the other hand, *rsad2* expression peaked on day 2 when normalized to WT levels (Fig 7B). Overall, these finding may suggest that *mk2b*-/- and *mk3-/-* induced higher interferon response during viral infection which might contribute further up regulation of interferon associated gene compared to WT control.

**Figure 7:**
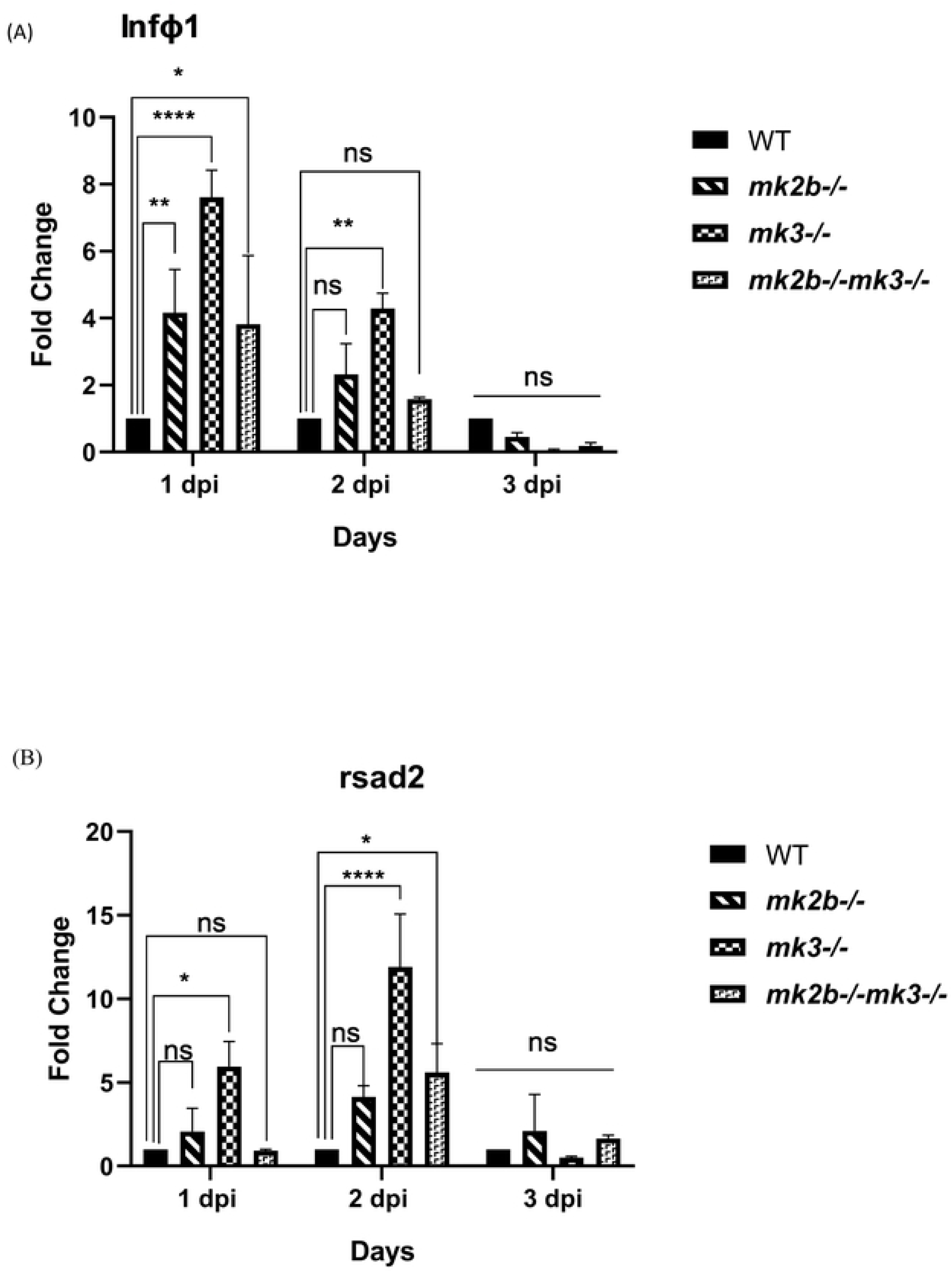
Expression levels of interferon (*infɸ1*) and interferon induced gene (*rsad2*) were upregulated after CHIKV infection: Total RNA was isolated from WT and knockout zebrafish larvae after infection and different genes were amplified by RT-qPCR. (A) Bar graph indicating the expression level of *infɸ1* and (B) *rsad2* in different knockout sets and WT. Data of three independent experiments are shown as mean ± SD. ****p<0.0001 was considered significant.

## Discussion

Host factors are essential at different stages of viral life cycle for efficient virion production and are therefore attractive targets for developing broad-spectrum antiviral therapeutics. Previously, we have reported that MK2 and MK3 are key host factors in CHIKV infection, using its inhibitors in cells and mice. In the current study, we examined the role of MK2 and MK3 during CHIKV infection at an organismal level by generating the single as well as double knockout of these two genes in zebrafish. This study incipiently explores and elucidate the role of MK2 and MK3 in a zebrafish knockout model.

Over seventy percent protein homology was seen of MK2 and MK3 between humans and zebrafish. To explore the role of *mk2b* and *mk3* during CHIKV infection, single knockouts of *mk2b* and *mk3*, along with a double knockout of *mk2b-mk3*, were successfully generated using the CRISPR-Cas9 technique. CHIKV titer were found to be significantly higher in all knockout groups compared to the WT control. Among the knockouts, the *mk3-/-* exhibited the highest viral titer post-infection, followed by *mk2b-/-*, while *mk2b-/-mk3-/-* double knockout showed the lowest viral titer. Furthermore, a large number of larvae showed severe symptoms (bent body, impaired response to physical stimuli and death) in the *mk3-/-* group. The expression levels of interferon (*infɸ1)* and interferon-induced gene (*rsad2)* were elevated across all knockout groups in the early days of infection, which might indicate elevated immune activation owing to high viral titer. Collectively, these results may highlight the essential roles of *mk2b* and *mk3* as key host factors in regulating CHIKV infection in organism level.

There are three major functions of MK2 and MK3 which include: cell cycle regulation, actin remodulation and post-transcriptional modification of TNF and other cytokines. Previous studies suggested that the knockout (KO) of MK2 leads to a decrease in proinflammatory cytokine production, which, in turn, makes the cells more susceptible to infection. For instance, knockout of MK2 in macrophage cell lines leads to decreased LPS-induced TNF (a proinflammatory cytokine) production (24). A similar result was observed in MK2 and MK3 knockout (KO) mouse models, where LPS exposure to MK2 KO mice resulted in a reduction of TNF level. In addition, the double knockout of MK2 and MK3 exhibited a striking alleviation in TNF level. On the other hand, overexpression of MK3 was found to compensate for the deficiency of MK2, suggesting that both the kinases share overlapping physiological functions *in vivo* but are expressed at different levels (25). Beyond LPS induction, studies have also examined bacterial infections in MK2 and MK3 knockout models. For instance, inhibition of p38MAPK or MK2 significantly increased bacterial loads in Salmonella-infected mouse embryonic fibroblasts (MEFs) and MK2-deficient (MK2 -/-) cells (25,26). Similarly, in case of *Listeria monocytogenes* infection, MK2-deficient mice exhibited heightened susceptibility. (26). This might be due to the significant reduction in TNF level, highlighting the critical role of MK2 in TNF induction during Gram-positive bacterial infection (25). This current study demonstrated that zebrafish with single or double knockouts of *mk2b* and *mk3* led to higher susceptibility to CHIKV infection compared to WT. Interferon (*infɸ1*) and interferon induced gene (*rsad2*) were also found to be upregulated in case of knockout larvae compared to WT control after CHIKV infection. Interferons regulate various MAPK pathways modulating the production of different proinflammatory cytokines which eventually act towards viral clearance (23). Additionally, MK2 KO has been shown to induce TNF response *in vivo* (25). Consequently, high viral titer as found in the *mk2b-/-* and *mk3-/-* compared to the WT control might result from the impaired TNF production. In addition to regulating TNF production, MK2 and MK3 are also involved in actin remodeling (12) and filopodia formation (24). Previous study from our group demonstrated that the presence of an MK2 inhibitor (CMPD1) prevents CHIKV particles from egressing out of the host cells. (12). In the current study, zebrafish were infected by CHIKV intravenously (CCV), which led to the dissemination of virus particles throughout the body and infected many cells. Preventing viral egress in absence of MK2 and MK3 might be another reason why knockout zebrafish exhibited higher viral titers compared to WT control. Collectively, these findings may suggest that impaired cytokine production and /or disrupted cytoskeletal dynamics in the absence of *mk2b* and *mk3* might lead to higher viral titer in knockout zebrafish as compared to WT control. However, further investigations are warranted towards extricating the mechanistic insights.

MK2 and MK3 are expressed and activated in parallel, however, MK2 showed significantly higher activity than MK3 (25). Conversely, other studies also reported disruptive roles of MK2 and MK3, where it has been depicted that the absence of MK3 alone aggravated the disease course in the absence of MK2 (28). Additionally, literature suggested that MK2 and MK3 regulate the expression of various mediators not only cooperatively but also through distinct mechanisms. Whole transcriptome analyses have identified many transcripts that are regulated uniquely in the absence of either MK2 or MK3, alongside the transcripts that are co-regulated (29). In zebrafish, it has been shown that the expression level of *mk3* is higher than that of *mk2b* (Sup Fig-3). In the present study, the high viral load observed in *mk3* knockout may indicate the possible regulation of certain transcripts by *mk3* independent of *mk2b* and/or higher expression levels of *mk3* in zebrafish. Regulation of different transcripts or activation of other downstream substrates or pathways can be monitored by the RNA sequencing in future.

The viral titer in case of double knockout *(mk2-/-mk3-/-)* was comparatively lower than that of single knockouts (*mk2b-/-* and *mk3-/-*). The possible attribute could be the activation of alternative innate immune signaling pathways immediately following infection. Other pathways that are reported to be activated after viral infection include: retinoic acid-inducible gene-I (RIG-I) (30) and melanoma differentiation-associated factor 5 (MDA5) pathways (31). These pathways are involved in activating downstream signaling proteins to generate an antiviral response in zebrafish. Subsequent future studies would be interesting to perceive more about the induction of other pathways in the absence of *mk2b* and *mk3*.

Regulation of CHIKV infection in the absence of *mk2b* and *mk3* has been presented in this study, however, exploring the mechanism and involvement of other pathways can be examined further. In conclusion, this study confirms that the host proteins *mk2b* and *mk3* are important players in controlling the CHIKV infection in organism level and subsequently can help in designing future antiviral therapeutics. Furthermore, knockout model of *mk2b* and *mk3* in zebrafish may provide a valuable tool for investigating their role in other viral infections.

## Materials and Methods

### Cell, virus and antibodies

Vero cells (African green monkey kidney cells) were obtained from the National Centre for Cell Science (NCCS), India and the CHIKV IS (accession no. PP349434) was used in this study. Cells were maintained in Dulbecco’s modified Eagle’s medium (DMEM, Himedia) supplemented with 10% Fetal Bovine Serum (FBS-Gibco), penicillin-streptomycin (Himedia) and gentamycin (Gibco). A monoclonal antibody of CHIKV-E2 was a kind gift from M.M. Parida (DRDE, Gwalior India) and Alexa flour-647 (Invitrogen) was used as secondary antibody.

### Bioinformatics analysis

The genome sequences and functions of *mk2a, mk2b* and *mk3* were obtained from the ZFIN server ( https://zfin.org). Alignment of the genes as well as proteins sequences between humans and zebrafish was performed using the Clustal Omega tool and the percent homology among the sequences was determined using NCBI blast (http://blast.ncbi.nlm.nih.gov).

### Ethics statement and zebrafish maintenance

AB strain of *Danio rerio* was used in the present study. All the experiments were conducted at the Institute of Life Sciences, Bhubaneswar with approval from the Institutional Animal Ethics Committee (IAEC) under proposal number: ILS/IAEC-250-AH/FEB-22. The embryos and larvae of zebrafish were raised in E3 media at 28.5°C. Adult zebrafish were maintained at a zebrafish maintaining system designed by Techniplast. The temperature and pH of the running water supply are maintained at 28.5°C and 7.5, respectively.

### Synthesis of gRNA

To generate the knockouts of *mk2b* and *mk3*, the CRISPR-Cas9 technique was used. Guide RNA (gRNA) sequences of 22 bases and 20 bases (including PAM sequence NGG) were selected using the CRISPR-RGEN tool to target the 5^th^ exon of both the *mk2b* and *mk3* genes (Table-1). The *mk2b* and *mk3* gRNAs were synthesized using the Invitrogen^TM^ MEGAScript^TM^ T7 transcription kit according to the procedure mentioned earlier (32).

### Generation of mk2b-/-, mk3-/- and mk2b-/-mk3-/-

To generate *mk2b-/-* and *mk3-/-, mk2b* and *mk3* gRNAs were microinjected separately into zebrafish embryos at the single-cell stage. For the generation of *mk2b-/-mk3-/-* double knockout, both the gRNAs of *mk2b* and *mk3* were microinjected into the same embryo. According to the protocol of Varshney et.al.(32) genotyping was performed from 8–10 microinjected zebrafish larvae 48 hours post-fertilization (hpf). Once the gRNA efficiency was confirmed, the remaining embryos were grown to adulthood as founder fish (F0). The positive F0 fishes were then backcrossed with wild-type fish to obtain the first filial generation (F1). Male and female F1 fishes with the same type of mutation were crossed to produce second filial generation fish (F2). After three months, F2 fishes were genotyped to identify homozygous mutants. Homozygous adult fishes with the same mutation were then transferred to a separate tank and embryos collected from their in-cross were used for further studies.

### High resolution melting curve analysis (HRM)

HRM is used to detect CRISPR-induced mutation in zebrafish during the process of generating knockouts. The detection process uses the advantage of a shift in the melt curve between WT and mutant target DNA. Firstly, RT-PCR was performed using the specific primers of *mk2b* and *mk3* (Table 1) which amplifies the region targeted by gRNA (gRNA against *mk2b* and *mk3*) followed by melt curve analysis. The genomic DNA, which was isolated from the zebrafish larvae and zebrafish fin for different experiments, was used as a template. The following run method was set for the amplification and the analysis of the shift of the melt curve: 95°C, 10 mins: (95°C, 15 secs: 60°C, 20 secs: 72°C, 20 secs) x 40; 95°C, 15 secs: 60°C, 15 secs: 95°C, 15 secs (33).

### Heteroduplex analysis (HD)

HD was performed by denaturing the amplified product (generated by HRMA) at 95°C for 5 mins, followed by reannealing at RT for 15 mins, which resulted in the formation of homoduplexes and heteroduplexes of complete complementary and incomplete complementary stands respectively. These products were then loaded onto 12% Polyacrylamide gel to check different homo and heteroduplex bands (33).

### CHIKV stock preparation and microinjection

The CHIKV IS (accession no. PP349434) was grown in Vero cells at a multiplicity of infection (MOI)-1 for 15 hrs. The cells and supernatant were then harvested. The cell-containing media was subjected to three freeze-thaw cycles and subsequently centrifuged. The titer of the collected virus was determined through plaque assay. Approximately 250 viral particles were used for microinjection into zebrafish larvae intravenously (30).

### Cloning and sequencing

To identify the mutation created on CRISPR target site, amplified products of *mk2b* and *mk3* genes were cloned into the pGEM®-T vector. These fragments were first amplified using genomic DNA as a template which was isolated from the F1 heterozygous mutant fishes. The amplified products were then cloned into the pGEM®-T Vector according to the manufacturer’s protocol, followed by transformation. The blue-white colony screening strategies were used to select positive colonies from the bacteria agar plate. White colonies were selected for plasmid isolation and sent for the Sanger DNA sequencing (31).

### Probe synthesis

To synthesize the probe for whole mount in situ hybridization (WISH) a part of *mk2b* and *mk3* genes were amplified using its specific primers. These amplified products were cloned in the PCR-Blunt II-TOPO vector (Invitrogen) followed by the linearization of these plasmids. The *mk2b* plasmid was linearized with BamHI and transcribed by T7 polymerase whereas the *mk3* plasmid was linearized with XhoI and transcribed by SP6 polymerase to synthesize the antisense RNA oligo. Digoxygenin (DIG) labelling mix (Roche 11277073910) was used to synthesize the labelled RNA probe (31).

### Plaque assay

To detect infectious virus particles, a total of 15 larvae were collected each day, up to 5 days post-infection (dpi). The collected larvae were homogenized in DMEM medium, followed by centrifugation, and the supernatant was collected for plaque assay. Vero cells were seeded in 12-well plates and infected with the collected samples. This assay was performed according to the procedure described previously (34). The cells were overlaid with methyl cellulose and incubated at 37°C for 3 days. The cells were then fixed with 8% formaldehyde and stained with crystal violet. Visible plaques were counted to determine viral titer.

### RT-qPCR

To assess CHIKV gene expression, approximately 15 larvae were collected daily from Day 1 to Day 5 post-infection and homogenized in RNA isolation reagent. RNA was isolated using the MagSure^®^ All-RNA isolation kit (RnaBio) followed by the synthesis of cDNA using the 1^st^ strand cDNA synthesis kit (TAKARA) as per the manufacturer’s instructions. An equal volume of cDNA was subjected to PCR amplification of the CHIKV E1 gene using its specific primers (Table 1) (35). Ef1α served as a control for gene expression.

### Whole mount in situ hybridization (WISH)

The whole-mount in situ hybridization (WISH) was performed using 2 dpi larvae. Briefly, approximately 20 larvae from infected sets of WT and knockouts were collected and fixed in 4% paraformaldehyde, followed by dehydration in methanol through a series. The samples were then stored at −20°C. The next day, hybridization was performed using the DIG labelled antisense RNA probe against the *mk2b* and *mk3* genes of zebrafish according to the protocol mentioned in Fatma et al., 2021 (33).

### Immunofluorescence

A total of 10 larvae were collected on 2 dpi and stored in 4% PFA. On the following day, the larvae were washed four times with 1X PBS-0.1% Triton-X, after which the solution was replaced with ice-cold acetone. The samples were then incubated at −20°C for 1 hr. Subsequently, the samples were subjected to blocking (1X PBS containing BSA, DMSO, and 10% Triton-X) for 2 hrs at room temperature, followed by the addition of the CHIKV-E2 primary antibody overnight.

Next day, the primary antibody was removed, and the samples were washed four times with 1X PBS-0.05% Tween. After washing, the wash buffer was replaced with Alexa Fluor 647-conjugated secondary antibody diluted in the blocking solution. The samples were incubated at room temperature for 1 hr and then transferred to 4°C for overnight incubation.

On the third day, the secondary antibody was removed, and the samples were washed before being refixed with 4% PFA for 20 minutes. The samples were then washed with 1X PBS-0.05% Tween, followed by mounting. The fluorescence intensity of CHIKV E2 protein was determined using a Thunder microscope (Leica M205 FA) (36).

### Statistical analysis

The statistical analysis was performed using the 1-way ANOVA and 2-way ANOVA method in graph pad prism 8.0.1 and the data was presented as mean ± SD. After analysis the data with 2-way ANOVA, the graphs represent the statistical significance where P<0.0001 is indicated by (****) and P=0.0001 is indicated by (**). Adjusted P value ranged between 0.0003-0.0008 is indicated by *** in the case of statistical analysis using 1-way ANOVA.

## Acknowledgments

We extend our sincere gratitude to Dr. M.M. Parida (DRDE, India) for providing the CHIKV-E2 antibody. We are also thankful to Kalyani Sahoo for her assistance with the ISH experiment. Additionally, we acknowledge Satyajit Behera and Laxmipriya Pattanaik for their support in zebrafish maintenance. We are grateful to Amrita Ray, Ankita Datey, Soumyajit Ghosh and Tathagata Mukherjee for their help especially during an urgent experiment and manuscript correction. My genuine appreciation goes to Mr Bhabani Shankar Sahoo from the confocal imaging facility. Finally, I extend my heartful appreciation to CSIR for their consistent financial support. SSK is a recipient of CSIR-SRF (113-2334-6704/2K23/1) and UN is a recipient of the DST-Inspire fellowship (IF180156). This work is also supported in part by Grant BT/PR42322/TRM/120/525/2021 from the Department of Biotechnology (DBT), Ministry of Science and Technology, Government of India and intramural funds from the BRIC-ILS, which is an autonomous institute of DBT, Government of India.

## Data availability

All the data generated have been included in the manuscript.

